# Associations of Mitochondrial Genomic Variation with Corticobasal Degeneration, Progressive Supranuclear Palsy, and Neuropathological Tau Measures

**DOI:** 10.1101/2020.07.10.196097

**Authors:** Rebecca R. Valentino, Nikoleta Tamvaka, Michael G. Heckman, Patrick W. Johnson, Alexandra I. Soto-Beasley, Ronald L. Walton, Shunsuke Koga, Ryan J. Uitti, Zbigniew K. Wszolek, Dennis W. Dickson, Owen A. Ross

## Abstract

Mitochondrial health is important in ageing and dysfunctional oxidative phosphorylation (OXPHOS) accelerates ageing and influences neurodegeneration. Mitochondrial DNA (mtDNA) codes for vital OXPHOS subunits and mtDNA background has been associated with neurodegeneration; however, no study has characterised mtDNA variation in Progressive supranuclear palsy (PSP) or Corticobasal degeneration (CBD) risk or pathogenesis. In this case-control study, 916 (42.5% male) neurologically-healthy controls, 1051 (54.1% male) pathologically-confirmed PSP cases, and 173 (51.4% male) pathologically-confirmed CBD cases were assessed to determine how stable mtDNA polymorphisms, in the form of mtDNA haplogroups, were associated with risk of PSP, risk of CBD, age of PSP onset, PSP disease duration, and neuropathological tau pathology measures for neurofibrillary tangles (NFT), neuropil threads (NT), tufted astrocytes (TA), and oligodendroglial coiled bodies (CB). 767 PSP cases and 152 CBD cases had quantitative tau pathology scores. mtDNA was genotyped for 39 unique SNPs using Agena Bioscience iPlex technologies and mitochondrial haplogroups were defined to mitochondrial phylogeny. After adjustment for multiple testing, we observed a significant association with risk of CBD for mtDNA sub-haplogroup H4 (OR=4.49, P=0.001) and the HV/HV0a haplogroup was associated with a decreased severity of NT tau pathology in PSP cases (P=0.0023). Our study reports that mitochondrial genomic background may be associated with risk of CBD and may be influencing tau pathology measures in PSP. Replication of these findings will be important.

## Introduction

Progressive supranuclear palsy (PSP) and Corticobasal degeneration (CBD) are rare progressive neurodegenerative movement disorders [1, 2]. PSP typically presents clinically with early falls, supranuclear vertical gaze palsy, parkinsonism, and dementia at about 65 years of age [3]. Individuals with CBD often present with progressive asymmetric rigidity and apraxia, loss of coordination, tremor, bradykinesia, akinesia, and occasionally alien limb syndrome [4, 5]. Both diseases have overlapping clinical symptoms with each other and other neurodegenerative diseases, such as Parkinson’s disease (PD) and Alzheimer’s disease (AD) [3, 6–8]. This can result in an inaccurate clinical diagnosis; definitive diagnosis is only achieved post-mortem using specific neuropathological diagnostic criteria [9, 10].

Neuropathologically, PSP and CBD are characterised as primary four-repeat (4R) tauopathies, with tau-positive aggregates in the form of neurofibrillary tangles (NFT), tufted astrocytes (TA), neuropil threads (NT), and oligodendroglia coiled bodies (CB), evident in the basal ganglia, diencephalon, and brainstem in PSP [11, 12], and in the substantia nigra and locus coeruleus in CBD [10]. Although generally considered sporadic disorders, *MAPT*, which encodes microtubule associated protein tau, is consistently documented as a strong genetic risk factor for both PSP and CBD [13, 14], and genetic variation in *MAPT* influences tau pathology severity in PSP [15]. Other genetic factors have also been identified however they do not explain complete disease aetiology [11, 13, 14, 16].

Age is the major risk factor for PSP and CBD, and mitochondrial health is well-established to contribute significantly to healthy ageing [17]. Mitochondrial dysfunction is also recognised in PSP pathogenesis as well as other clinically similar diseases such as PD and AD [18–21]. More specifically, defective mitochondria generate reactive oxygen species (ROS) which oxidise proteins, lipids, and nucleic acids, accelerating the ageing process [22]. ROS is suggested to contribute to the accumulation of insoluble proteinaceous deposits, such as Lewy bodies in PD, and senile plaques and NFT in AD [23–25], and dysfunction of complex I in the oxidative phosphorylation (OXPHOS) system has been shown to accelerate 4R tau isoform formation in PSP cell lines [26] and is defective in the substantia nigra of PD patients [27, 28].

Mitochondria contain their own double-stranded, circular 16.6 kilo-base pair genome (mtDNA), independent to nuclear DNA (nDNA). mtDNA encodes 37 polypeptides, of which 13 encode vital OXPHOS subunits. An individual cell can contain hundreds to thousands of mtDNA copies which significantly affects cellular metabolic background [29]. mtDNA also contains stable single nucleotide polymorphisms which define individuals to specific haplogroups. Individual mtDNA haplogroups have distinctive metabolic demands [30, 31] and haplogroup bioefficiency has also been shown to affect ageing and risk of developing many neurodegenerative diseases, including PD and AD [31–34].

Despite evidence reporting mitochondrial dysfunction in PSP, no studies have examined if mtDNA background influences PSP or CBD risk or if mtDNA variation can contribute to overall tau pathology severity. Thus, herein we examine the association of mtDNA background with PSP and CBD risk and tau pathology severity in two autopsy-defined series.

## Methods

### Study Design

1051 pathologically confirmed PSP cases, 173 pathologically confirmed CBD cases, and 916 neurologically healthy controls were included. All subjects were of self-reported European descent. PSP samples were donated between 1998 and 2016 and CBD samples were collected between 1994 and 2017. All samples were obtained from the CurePSP Brain Bank at Mayo Clinic Jacksonville and were rendered by a single neuropathologist (DWD) following published criteria [8, 10, 35]. Controls were recruited from 1998 to 2015 through the clinical Neurology department at Mayo Clinic Jacksonville, Florida. Demographic information is summarised in Table 1. Age of onset and disease duration was unavailable for 611 PSP and was not available for CBD cases. This study was approved by the Mayo Clinic Institutional Review Board and individual written consent was obtained from all subjects, or their next of kin, prior to commencement.

**Table 1:** Summary of cohort characteristics in N=1051 PSP cases, 173 CBD cases, and N=916 controls. The sample median (minimum, maximum) is given for continuous variables. CB=coiled bodies; NFT=neurofibrillary tangles; TA=tufted astrocytes; NT=neuropil threads.

| Variable | PSP cases (N=1051) | CBD cases (N=173) | Controls (N=916) |
| --- | --- | --- | --- |
| Age (years) | 75 (45, 98) | 70 (46, 96) | 79 (41, 102) |
| Sex |  |  |  |
| Male | 569 (54.1%) | 89 (51.4%) | 389 (42.5%) |
| Female | 482 (45.9%) | 84 (48.6%) | 527 (57.5%) |
| Age of onset (years) | 68 (41, 90) | -- | -- |
| Disease duration (years) | 7 (1, 32) | -- | -- |
| PSP clinical subtype |  |  |  |
| Richardson | 570 (74.4%) | -- | -- |
| Non-Richardson | 196 (25.6%) | -- | -- |
| Braak stage |  |  |  |
| 0 | 113 (14.7%) | 20 (13.2%) | -- |
| I | 128 (16.7%) | 33 (21.7%) | -- |
| II | 224 (29.2%) | 51 (33.6%) | -- |
| III | 235 (30.6%) | 39 (25.7%) | -- |
| IV | 50 (6.5%) | 7 (4.6%) | -- |
| V | 11 (1.4%) | 1 (0.7%) | -- |
| VI | 6 (0.8%) | 1 (0.7%) | -- |
| Thal phase |  |  |  |
| 0 | 337 (43.9%) | 83 (54.6%) | -- |
| 1 | 126 (16.4%) | 31 (20.4%) | -- |
| 2 | 53 (6.9%) | 14 (9.2%) | -- |
| 3 | 188 (24.5%) | 19 (12.5%) | -- |
| 4 | 43 (5.6%) | 3 (2.0%) | -- |
| 5 | 20 (2.6%) | 2 (1.3%) | -- |
| CB tau pathology score | 1.50 (0.25, 2.36) | 0.76 (0.23, 1.75) | -- |
| NFT tau pathology score | 2.23 (0.83, 2.89) | 2.19 (0.99, 2.67) | -- |
| TA tau pathology score | 1.00 (0.06, 2.00) | 0.52 (0.24, 1.04) | -- |
| NT tau pathology score | 2.15 (0.35, 2.90) | 2.52 (1.23, 2.95) | -- |

### Neuropathological Assessment

Semi-quantitative tau pathology scores in PSP and CBD cases were determined by a single neuropathologist (DWD) using standardized histopathologic methods and phospho-tau immunochemistry. Scores were generated in a subset of cases (PSP: N=767, CBD: N=152) on a four-point severity scale (0=none, 1=mild, 2=moderate, and 3=severe) [36]. All sections from all cases were processed in an identical manner with phospho-tau monoclonal antibody (CP13, from Dr. Peter Davies, Feinstein Institute, Long Island, NY) and immunohistochemistry using a DAKO Autostainer. NFT (PSP: N=767, CBD: N=152), CB (PSP: N=766, CBD: N=152), TA (PSP: N=737, CBD: N=152), and NT (PSP: N=766, CBD: N=151) were assessed, and overall scores in 17–20 different, vulnerable neuroanatomical regions in PSP/CBD were generated for each separate tau pathology measure (Supplementary Table 1). Mean semi-quantitative measures were then calculated for each PSP/CBD patient across all anatomical regions, where a higher overall score indicated more severe tau pathology. PSP/CBD patients who did not have tau scores in a given region for a given tau pathology measure had their scores imputed by using the mean of the values of the patients in the given disease group (PSP or CBD) who did have scores. Any patients with missing data for >50% of neuroanatomical regions for a given tau pathology measure were not included in any analysis involving tau pathology measures. PSP and CBD cases were additionally assessed for Alzheimer-type pathology with thioflavin-S fluorescent microscopy. Braak NFT stage [37] and Thal amyloid phase [38] were generated for each case based on the density and distribution of plaques and tangles, as previously detailed [39, 40] (Table 1).

### DNA Preparation and Genotyping

Genomic DNA and mtDNA was extracted from peripheral blood lymphocytes from control subjects and from frozen cerebellum brain tissue from PSP and CBD cases using Autogen Flex Star and Autogen 245T (Holliston, MA) methods respectively. mtDNA in all samples was genotyped on two high multiplex custom-designed iPlex assays (consisting of 39 mtDNA SNPs - Figure 1) using Sequenom MassARRAY iPlex technology (MALDI-TOF MS) and iPlex^®^ Gold chemistry technology [41]. PCR and sequence-specific extension primers were designed through Sequenom’s MassARRAY Typer 4.0 Designer software (version 0.2) (Supplementary Table 2), part of the Assay Design Suite (Agena Bioscience™, San Diego, USA), and were manufactured by Integrated DNA Technologies (IDT, San Diego, USA). Genotypes were determined by Sequenom MassARRAY^®^ Bruker mass spectrometry (Agena Bioscience, San Diego, CA, USA) [41] and were accepted if intensities were >5 from the base line intensity (<5 was considered noise). Individuals with a genotyping call rate >95% were accepted and had mtDNA haplogroups determined. Genotyping analysis was conducted using Sequenom’s Typer 4.0 Analyzer software (version 25.73). Additional details are provided in Supplementary Methods.

**Figure 1:**
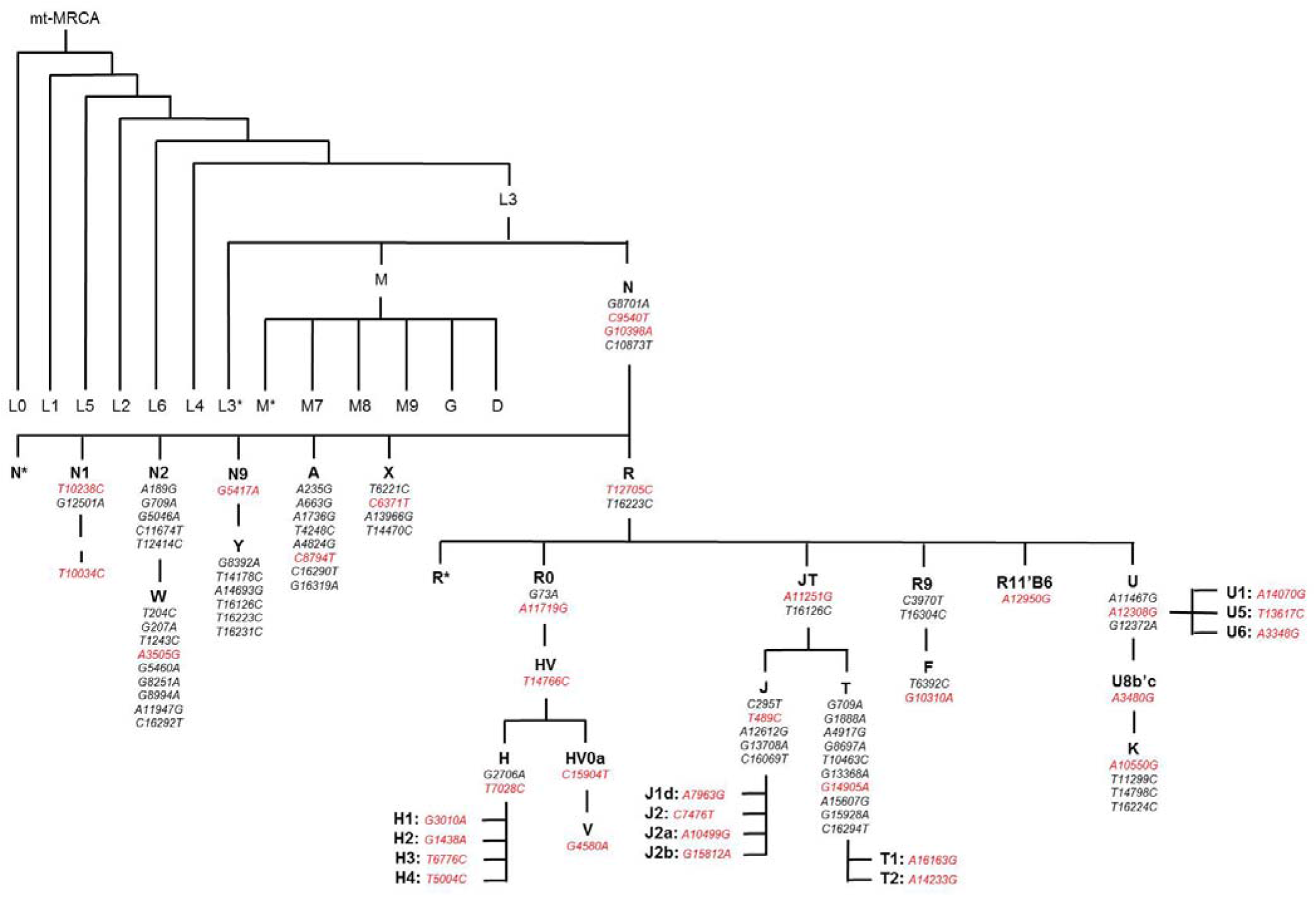
Schematic overview of the mitochondrial phylotree and unique SNPs which define European mitochondrial haplogroups. Mitochondrial SNPs highlighted in red indicate SNPs genotyped using Sequenom iPlex technology (Agena Bioscience, San Diego, CA, USA) to determine mitochondrial DNA haplogroups. Adapted from Phylotree^1^.

### Mitochondrial DNA Haplogroup Assignment

Mitochondrial DNA haplogroups were manually defined to mitochondrial phylogeny [42, 43] (Figure 1). Haplogroups and sub-haplogroups were determined whereby patterns of mtDNA SNPs had to sequentially be present down the phylotree and not present in other phylogenetic clades (refer to Supplementary Methods for further details). For use in secondary analysis, super-haplogroups were determined by combining phylogenetic-related haplogroups together (Supplementary Methods). Over 95% of individuals in European populations classify as one of the following mtDNA haplogroups - H, V, J, T, I, X, W, U, or K [44], therefore individuals with a non-European mitochondrial haplogroups (e.g. non-N, A, F, and B) were removed from analysis.

### Statistical Analysis

Associations of mitochondrial haplogroups with risk of PSP and CBD (each separately versus controls) were evaluated using logistic regression models that were adjusted for age and sex. Odds ratios (ORs) and 95% confidence intervals (CIs) were estimated. In analysis of only PSP or CBD patients, linear regression models were used to assess associations of mitochondrial haplogroups with PSP disease duration, age of PSP onset, and tau pathology scores of; CB, NFT, TA, and NT. Models were adjusted for age of PSP onset and sex (models involving PSP disease duration), for sex (models involving age of PSP onset), and for age at death, sex, Braak stage, and Thal phase (models involving CB, NFT, TA, and NT tau pathology scores).

Haplogroups that occurred in <10 subjects for a given association analysis were not included in that analysis. For the primary analysis (i.e. all analysis not involving super-haplogroups), a Bonferroni correction was applied for multiple testing separately for each group of similar statistical tests. Specifically, p-values ≤0.0019 (associations with PSP), ≤0.0023 (associations with CBD), ≤0.0025 (associations with disease duration and age of PSP onset), ≤0.0024 and ≤0.0045 (associations with tau pathology scores in either PSP or CBD cases respectively) were considered to be statistically significant. No adjustment for multiple testing was made for secondary analysis involving super-haplogroups and p-values ≤0.05 were considered significant. A power analysis regarding associations of mitochondrial haplogroups with disease risk and tau pathology scores is displayed in Supplementary Table 3. Statistical analyses were performed using R Statistical Software (version 3.6.1).

## Results

Mitochondrial DNA haplogroup frequencies in our control cohort were representative of European populations (Table 2) [43] and were considered an appropriate reference cohort to investigate mtDNA background associations with risk of PSP and CBD. Individuals carrying non-European mtDNA haplogroups (non-N, A, F, and B) were previously removed to ensure individuals from European descent were assessed. In analysis that was adjusted for age and sex, to remove possible confounding influences, and after correcting for multiple testing (P≤0.0019 considered significant), there were no significant associations between individual mtDNA haplogroups and PSP risk (all P≥0.040, Table 2). We did observe a significant association between mtDNA haplogroup H4 and an increased risk of CBD (5.2% vs. 1.2%, OR=4.49, P=0.001, Table 2).

**Table 2:** Associations of individual mitochondrial DNA haplogroups with risk of PSP and CBD (compared to controls) were evaluated using multivariable logistic regression models adjusted for age and sex. After applying a Bonferroni correction for multiple testing, association p-values≤0.0019 (PSP vs. controls analysis) and ≤0.0023 (CBD vs. controls) are considered statistically significant. ^1^Statistical tests were not performed for these haplogroups owing to their rare frequency (<10 subjects in the given haplogroup for the given comparison [PSP vs. controls or CBD vs. controls]). OR=odds ratio; CI=confidence interval at 95%.

| Mitochondrial DNA Haplogroup | Haplogroup frequency, No. (%) |  |  | PSP vs. controls |  | CBD vs. controls |  |
| --- | --- | --- | --- | --- | --- | --- | --- |
|  | Controls (N=916) | PSP patients (N=1051) | CBD patients (N=173) | OR (95% CI) | P-value | OR (95% CI) | P-value |
| N<sup>1</sup> | 2 (0.2%) | 0 (0.0%) | 0 (0.0%) | -- | -- | -- | -- |
| N1 | 5 (0.5%) | 6 (0.6%) | 0 (0.0%) | 1.08 (0.32, 3.79) | 0.90 | -- | -- |
| I | 31 (3.4%) | 23 (2.2%) | 2 (1.2%) | 0.56 (0.32, 0.97) | 0.040 | 0.24 (0.04, 0.83) | 0.056 |
| W | 15 (1.6%) | 22 (2.1%) | 3 (1.7%) | 1.37 (0.70, 2.72) | 0.36 | 1.26 (0.29, 4.00) | 0.72 |
| X | 8 (0.9%) | 18 (1.7%) | 5 (2.9%) | 1.89 (0.84, 4.67) | 0.14 | 3.47 (0.99, 11.09) | 0.039 |
| R and R0<sup>1</sup> | 6 (0.7%) | 10 (1.0%) | 2 (1.2%) | 1.33 (0.49, 3.96) | 0.59 | -- | -- |
| HV and HV0a | 22 (2.4%) | 16 (1.5%) | 2 (1.2%) | 0.67 (0.34, 1.29) | 0.23 | 0.53 (0.08, 1.88) | 0.40 |
| H, H1, H2, H3, and H4 | 423 (46.2%) | 469 (44.6%) | 84 (48.6%) | 0.94 (0.78, 1.12) | 0.48 | 1.10 (0.79, 1.53) | 0.59 |
| H | 199 (21.7%) | 200 (19.0%) | 30 (17.3%) | 0.84 (0.67, 1.05) | 0.13 | 0.75 (0.48, 1.14) | 0.19 |
| H1 | 145 (15.8%) | 171 (16.3%) | 31 (17.9%) | 1.02 (0.80, 1.31) | 0.85 | 1.15 (0.73, 1.76) | 0.54 |
| H2 | 36 (3.9%) | 34 (3.2%) | 5 (2.9%) | 0.78 (0.48, 1.26) | 0.31 | 0.65 (0.22, 1.58) | 0.38 |
| H3 | 32 (3.5%) | 49 (4.7%) | 9 (5.2%) | 1.47 (0.93, 2.35) | 0.10 | 1.75 (0.76, 3.67) | 0.16 |
| H4 | 11 (1.2%) | 15 (1.4%) | 9 (5.2%) | 1.18 (0.54, 2.66) | 0.68 | 4.49 (1.75, 11.27) | 0.001 |
| V | 18 (2.0%) | 29 (2.8%) | 3 (1.7%) | 1.36 (0.75, 2.53) | 0.31 | 0.79 (0.18, 2.43) | 0.71 |
| JT<sup>1</sup> | 2 (0.2%) | 3 (0.3%) | 0 (0.0%) | -- | -- | -- | -- |
| J, J1, J1d, J2a, and J2b | 93 (10.2%) | 125 (11.9%) | 18 (10.4%) | 1.23 (0.92, 1.64) | 0.16 | 1.11 (0.62, 1.87) | 0.71 |
| J<sup>1</sup> | 0 (0.0%) | 0 (0.0%) | 0 (0.0%) | -- | -- | -- | -- |
| J1 | 72 (7.9%) | 98 (9.3%) | 17 (9.8%) | 1.22 (0.89, 1.69) | 0.22 | 1.34 (0.74, 2.33) | 0.31 |
| J1d<sup>1</sup> | 1 (0.1%) | 0 (0.0%) | 0 (0.0%) | -- | -- | -- | -- |
| J2a | 13 (1.4%) | 24 (2.3%) | 1 (0.6%) | 1.77 (0.90, 3.62) | 0.10 | 0.48 (0.03, 2.54) | 0.49 |
| J2b | 7 (0.8%) | 3 (0.3%) | 0 (0.0%) | 0.39 (0.08, 1.41) | 0.17 | -- | -- |
| T, T1, and T2 | 77 (8.4%) | 102 (9.7%) | 22 (12.7%) | 1.20 (0.88, 1.64) | 0.26 | 1.71 (0.99, 2.83) | 0.044 |
| T<sup>1</sup> | 0 (0.0%) | 2 (0.2%) | 1 (0.6%) | -- | -- | -- | -- |
| T1 | 17 (1.9%) | 22 (2.1%) | 4 (2.3%) | 1.10 (0.58, 2.12) | 0.78 | 1.15 (0.32, 3.28) | 0.80 |
| T2 | 60 (6.6%) | 78 (7.4%) | 17 (9.8%) | 1.19 (0.84, 1.70) | 0.33 | 1.73 (0.94, 3.03) | 0.066 |
| U, U1, U3, U5, and U6 | 130 (14.2%) | 138 (13.1%) | 19 (11.0%) | 0.88 (0.68, 1.15) | 0.35 | 0.70 (0.40, 1.16) | 0.18 |
| U | 44 (4.8%) | 56 (5.3%) | 7 (4.0%) | 1.07 (0.71, 1.61) | 0.75 | 0.83 (0.33, 1.80) | 0.66 |
| U1<sup>1</sup> | 1 (0.1%) | 1 (0.1%) | 0 (0.0%) | -- | -- | -- | -- |
| U3<sup>1</sup> | 8 (0.9%) | 0 (0.0%) | 0 (0.0%) | -- | -- | -- | -- |
| U5 | 74 (8.1%) | 80 (7.6%) | 12 (6.9%) | 0.91 (0.65, 1.28) | 0.59 | 0.79 (0.39, 1.45) | 0.47 |
| U6<sup>1</sup> | 3 (0.3%) | 1 (0.1%) | 0 (0.0%) | -- | -- | -- | -- |
| K | 78 (8.5%) | 80 (7.6%) | 11 (6.4%) | 0.92 (0.66, 1.28) | 0.62 | 0.72 (0.35, 1.35) | 0.34 |

Associations of individual mtDNA haplogroups with tau pathology scores of CB, NFT, TA, and NT in PSP and CBD are summarised in Tables 3 and 4, respectively. After correction for multiple testing (P≤0.0023 considered significant) and when adjusting for age at death, sex, Braak stage, and Thal phase, PSP individuals with a haplogroup HV and HV0a background (N=10) had significantly lower NT pathology (P=0.0023, Table 3 and Supplementary Table 5) compared to other individuals; mean NT tau pathology scores were 0.35 units lower for haplogroup HV and HV0a cases (Supplementary Figure 1A). Additionally, although not quite statistically significant, mean NFT tau pathology score was 0.09 units lower for individuals with PSP and mtDNA haplogroup T, T1, and T2 backgrounds (N=85) compared to other individuals (P=0.009, Supplementary Figure 1B). No individual mtDNA haplogroups were significantly associated with any tau pathology measures in CBD (Table 4), however in secondary analysis super-haplogroup UK reported a lower CB tau pathology (Supplementary Table 6, P=0.013).

**Table 3:**
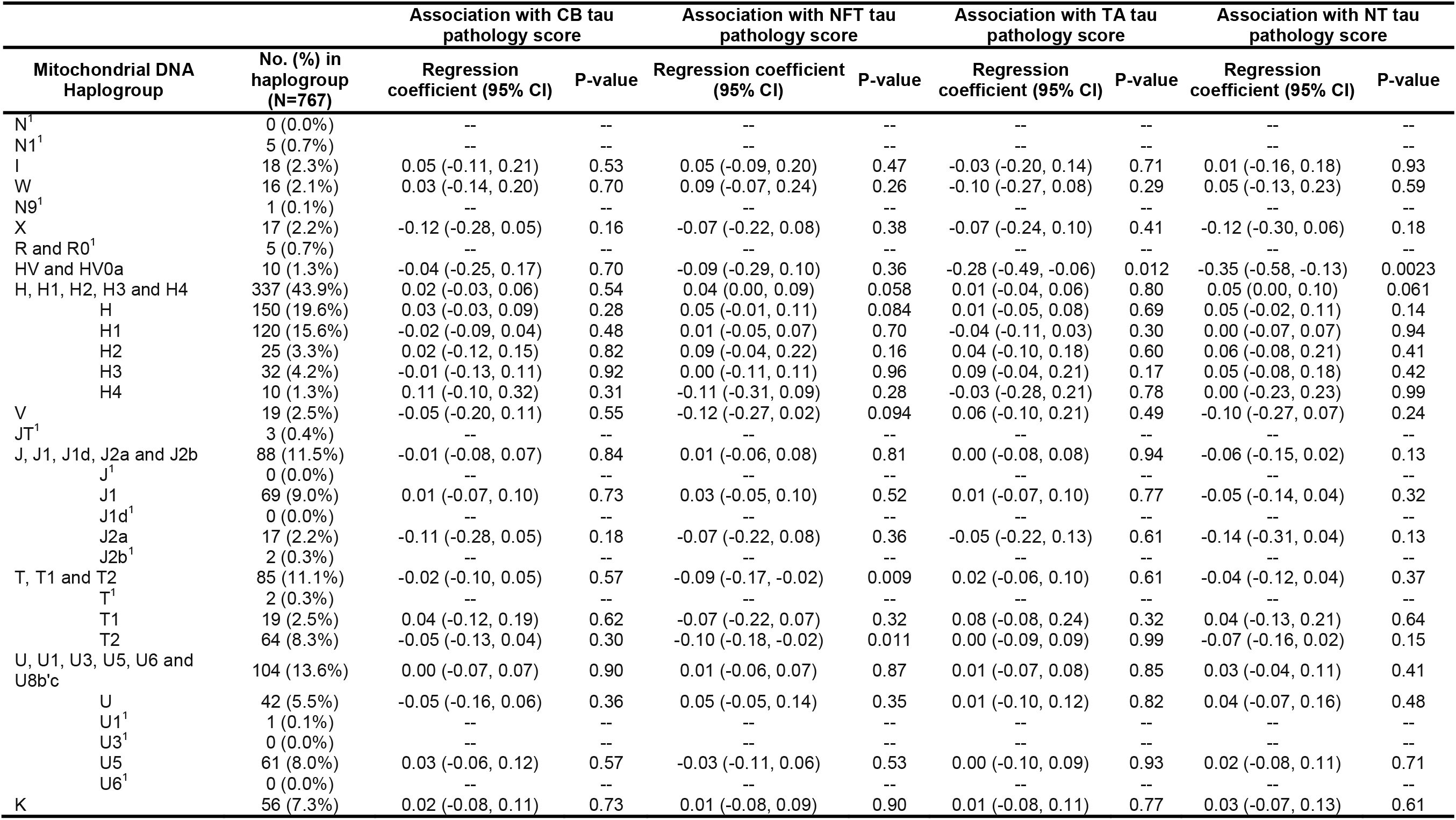
Associations of individual mitochondrial DNA haplogroups with CB, NFT, TA, and NT tau pathology scores in PSP cases with measured tau pathology scores (N=767) from linear regression models that were adjusted for age at death, sex, Braak, and Thal phase. Regression coefficients are interpreted as the increase in mean CB, NFT, TA, or NT tau pathology scores for patients in the given mitochondrial DNA haplogroup compared to patients not in the given haplogroup (non-haplogroup). P-values ≤0.0024 are considered statistically significant after applying a Bonferroni correction for multiple testing. ^1^Statistical tests were not performed for these haplogroups owing to their rare frequency (<10 PSP cases in the given haplogroup). CB=coiled bodies; NFT=neurofibrillary tangles; TA=tufted astrocytes; NT=neuropil threads; CI=confidence interval.

**Table 4:** Associations of individual mitochondrial DNA haplogroups with CB, NFT, TA, and NT tau pathology scores in CBD cases with measured tau pathology scores (N=152) from linear regression models that were adjusted for age at death, sex, Braak, and Thal phase. Regression coefficients are interpreted as the increase in mean CB, NFT, TA, or NT tau pathology scores for patients in the given mitochondrial DNA haplogroup compared to patients not in the given haplogroup (non-haplogroup). P-values ≤0.0045 are considered statistically significant after applying a Bonferroni correction for multiple testing. ^1^Statistical tests were not performed for these haplogroups owing to their rare frequency (<10 PSP cases in the given haplogroup, with the exception of haplogroup H4 which was examined despite the fact that it occurred in only 8 cases owing to the fact that it was of specific interest due to its significant association with risk of CBD). CB=coiled bodies; NFT=neurofibrillary tangles; TA=tufted astrocytes; NT=neuropil threads; CI=confidence interval.

| Mitochondrial DNA Haplogroup | No. (%) in haplogroup (N=152) | Association with CB tau pathology score |  | Association with NFT tau pathology score |  | Association with TA tau pathology score |  | Association with NT tau pathology score |  |
| --- | --- | --- | --- | --- | --- | --- | --- | --- | --- |
|  |  | Regression coefficient (95% CI) | P-Value | Regression coefficient (95% CI) | P-Value | Regression coefficient (95% CI) | P-Value | Regression coefficient (95% CI) | P-Value |
| I<sup>1</sup> | 1 (0.7%) | -- | -- | -- | -- | -- | -- | -- | -- |
| W<sup>1</sup> | 3 (2.0%) | -- | -- | -- | -- | -- | -- | -- | -- |
| X<sup>1</sup> | 4 (2.6%) | -- | -- | -- | -- | -- | -- | -- | -- |
| R and R0<sup>1</sup> | 2 (1.3%) | -- | -- | -- | -- | -- | -- | -- | -- |
| HV and HV0a<sup>1</sup> | 2 (1.3%) | -- | -- | -- | -- | -- | -- | -- | -- |
| H, H1, H2, H3 and H4 | 71 (46.7%) | 0.02 (-0.07, 0.11) | 0.70 | -0.01 (-0.09, 0.08) | 0.87 | 0.00 (-0.04, 0.04) | 0.96 | -0.02 (-0.12, 0.07) | 0.64 |
| H | 25 (16.4%) | 0.03 (-0.09, 0.15) | 0.67 | 0.12 (0.01, 0.23) | 0.035 | 0.01 (-0.05, 0.06) | 0.84 | 0.10 (-0.03, 0.22) | 0.13 |
| H1 | 26 (17.1%) | 0.05 (-0.07, 0.16) | 0.46 | -0.08 (-0.19, 0.04) | 0.19 | 0.00 (-0.06, 0.06) | 0.93 | -0.03 (-0.16, 0.10) | 0.61 |
| H2<sup>1</sup> | 4 (2.6%) | -- | -- | -- | -- | -- | -- | -- | -- |
| H3<sup>1</sup> | 8 (5.3%) | -- | -- | -- | -- | -- | -- | -- | -- |
| H4 | 8 (5.3%) | -0.09 (-0.28, 0.11) | 0.40 | 0.12 (-0.07, 0.31) | 0.21 | 0.04 (-0.06, 0.13) | 0.44 | 0.10 (-0.11, 0.31) | 0.34 |
| V<sup>1</sup> | 3 (2.0%) | -- | -- | -- | -- | -- | -- | -- | -- |
| J1 and J2a | 14 (9.2%) | 0.09 (-0.06, 0.25) | 0.23 | -0.01 (-0.15, 0.14) | 0.94 | 0.02 (-0.06, 0.09) | 0.67 | 0.09 (-0.07, 0.26) | 0.25 |
| J1 | 13 (8.6%) | 0.07 (-0.09, 0.23) | 0.40 | -0.01 (-0.16, 0.15) | 0.94 | -0.01 (-0.09, 0.06) | 0.71 | 0.08 (-0.09, 0.25) | 0.34 |
| J2a<sup>1</sup> | 1 (0.7%) | -- | -- | -- | -- | -- | -- | -- | -- |
| T, T1 and T2 | 21 (13.8%) | 0.08 (-0.05, 0.21) | 0.24 | 0.04 (-0.08, 0.16) | 0.53 | 0.02 (-0.04, 0.08) | 0.48 | 0.01 (-0.12, 0.15) | 0.83 |
| T<sup>1</sup> | 1 (0.7%) | -- | -- | -- | -- | -- | -- | -- | -- |
| T1<sup>1</sup> | 4 (2.6%) | -- | -- | -- | -- | -- | -- | -- | -- |
| T2 | 16 (10.5%) | 0.09 (-0.05, 0.24) | 0.20 | 0.02 (-0.11, 0.16) | 0.73 | 0.01 (-0.06, 0.08) | 0.74 | 0.01 (-0.14, 0.16) | 0.90 |
| U and U5 | 18 (11.8%) | -0.11 (-0.24, 0.03) | 0.12 | 0.02 (-0.11, 0.15) | 0.73 | -0.05 (-0.12, 0.02) | 0.14 | -0.05 (-0.20, 0.10) | 0.50 |
| U<sup>1</sup> | 7 (4.6%) | -- | -- | -- | -- | -- | -- | -- | -- |
| U5 | 11 (7.2%) | 0.02 (-0.15, 0.20) | 0.78 | 0.01 (-0.15, 0.18) | 0.89 | -0.07 (-0.15, 0.01) | 0.10 | 0.03 (-0.15, 0.21) | 0.73 |
| K | 11 (7.2%) | -0.16 (-0.33, 0.01) | 0.072 | -0.05 (-0.21, 0.12) | 0.58 | -0.01 (-0.09, 0.07) | 0.82 | -0.03 (-0.22, 0.15) | 0.73 |

Mitochondrial DNA background was not significantly associated with age of onset or disease duration in PSP (Supplementary Table 7); however, there was a suggestive association between mtDNA haplogroup W (N=11) and a longer disease duration in PSP (P=0.003, Supplementary Figure 1C). Key findings of our study are summarised in Supplementary Figure 2.

## Discussion

Mitochondrial health plays a significant role in ageing and the development of neurodegenerative diseases, including tauopathy [17–21]; however population specific mtDNA variation has not been investigated in PSP and CBD. Our findings indicate that major mtDNA haplogroups do not associate with risk of PSP; however, individuals with mtDNA sub-haplogroup H4 background may be at a significantly increased risk of CBD. In PSP cases, individuals with mtDNA haplogroup HV/HV0a backgrounds had a significantly decreased NT tau pathology and individuals with a haplogroup T (including T1 and T2) background had mildly reduced NFT tau pathology levels. No mtDNA haplogroups were significantly associated with tau pathology severity in CBD cases.

mtDNA sub-haplogroup H4, which was associated with an increased risk of CBD, is defined by a synonymous coding variant rs41419549 which is located in the NADH dehydrogenase subunit-2 (*MT-ND2*) (Supplementary Figure 2). ND2 has been identified to play a central role in the assembly of complex I subunits [45]. CBD pathology is localised to the substantia nigra and locus coeruleus [9], which may indicate mtDNA haplogroup H4 has a tissue specific effect on mitochondrial functionality. Albeit interesting, the absence of a H4 association with any tau pathology severity measure in CBD suggests that the presence of haplogroup H4 may be accelerating degeneration rather than enhancing tau aggregation. Despite the strong effect size (OR=4.49) and the fact that this association survived a stringent Bonferroni correction for multiple testing, as the H4 mtDNA haplogroup occurred in a small number of CBD patients (N=9), validation of this finding will be important.

Regarding mtDNA haplogroup HV/HV0a, which was associated with less NT tau pathology in PSP cases, haplogroup HV is defined by a single missense variant rs193302980 in the cytochrome-b subunit of complex III (*MT-CYB*) and haplogroup HV0a is defined by a unique variant rs35788393 located adjacent to *MT-CYB* in the coding region for tRNA-threonine (Supplementary Figure 2). Cytochrome-b is a vital component of complex III and ND6 is a subunit of complex I. Both complex I and III are important components of the OXPHOS pathway, and have been identified as major drivers of neurodegeneration and dysfunction of complex I has also been shown to accelerate 4R tau isoform formation in PSP cell lines [26, 27]. Again, despite the statistically significant association between mtDNA haplogroup HV/HV0a background and NT severity, replication of this finding will be key given that there were only 10 HV/HV0a PSP cases.

Although this is the first study to report significant associations of mtDNA variation background with CBD risk and with tau pathology in PSP, several limitations need to be acknowledged. First, to the best of our knowledge, all participants in our study were of European descent based on both self-reported ethnicity as well as mtDNA haplogroup profile (non-European haplogroups were excluded). Nonetheless, given the absence of available genome-wide population control markers, which would have allowed us to adjust our regression models for genetic background (in the form of top principal components from genetic data), we cannot rule out the possibility that population stratification could have had an effect on our results.

Another limitation is the lack of a replication series. Although large numbers of PSP and CBD cases were included in this study, considering the rarity of these two diseases, independent validation of our findings, as well as meta-analytic studies, will be important. Furthermore, power to detect associations was limited in the smaller CBD series and for rare haplogroups in both series.

Given the complex nature of mtDNA variation, levels of heteroplasmy may also be a concern in brain tissue. We used a PCR amplification-based MALDI-TOF MS technology which is considered sensitive enough to accurately determine alleles from pools of recombinants and is thus suitable for mtDNA-based population studies, limiting the impact of heteroplasmy and determining the individual mtDNA background [46]. Furthermore, the cerebellum is unaffected in PSP and CBD pathology and white blood cells are renewed every few days, therefore heteroplasmy levels are assumed to be low in our cases and controls and should not interfere with genotyping results in this study. In the future, performing mtDNA sequencing may identify higher impact and rarer penetrant variants. African and Asian haplogroup clades were also not investigated in this study and would need to be explored in future work. Finally, functional studies need to be performed to better understand the mechanisms by which mtDNA haplogroup background is contributing to disease risk and tau pathology severity.

## Conclusions

This is the first study to characterise the role of mtDNA background in susceptibility to PSP and CBD and in tau pathology severity in general. Our data suggests that mtDNA haplogroup background is associated with CBD risk and may also modify tau aggregation formation in PSP. Though larger validation studies will be key (particularly for CBD due to the smaller sample size of this group), it will also be important for future studies to investigate how established nDNA risk factors, such as the *MAPT* H1 haplotype, interact with mtDNA genetic background with regard to susceptibility to disease and severity of tau pathology.

## List of abbreviations

PSP: Progressive supranuclear palsy
CBD: Corticobasal degeneration
NFT: Neurofibrillary tangles
NT: Neuropil threads
TA: Tufted astrocytes
CB: Oligodendroglial coiled bodies
PD: Parkinson’s disease
AD: Alzheimer’s disease
4R: Four-repeat tau isoform
ROS: Reactive oxygen species
OXPHOS: Oxidative phosphorylation
mtDNA: Mitochondrial DNA
nDNA: Nuclear DNA
MALDI-TOF MS: Matrix-assisted laser desorption/ionization mass spectrometry
OR: Odds ratio
CI: Confidence interval
MT-ND2: NADH dehydrogenase subunit-2
MT-CYB: Cytochrome-b subunit of complex III
mtSNP: Mitochondrial DNA single nucleotide polymorphism

## Declarations

### Ethics approval

This study was approved by the Mayo Clinic Institutional Review Board and individual written consent was obtained from all participants, or their next of kin, prior to commencement.

### Consent for publication

Not applicable

### Availability of data and material

The datasets generated and/or analysed during the current study are available from the corresponding author on reasonable request.

### Competing interests

ZKW serves as PI or Co-PI on Abbvie, Inc. (M15-562 and M15-563), Biogen, Inc. (228PD201) grant, and Biohaven Pharmaceuticals, Inc. (BHV4157-206 and BHV3241-301). He serves as PI of the Mayo Clinic American Parkinson Disease Association (APDA) Information and Referral Center, and as Co-PI of the Mayo Clinic APDA Center for Advanced Research. All other authors declare that they have no competing interests.

### Funding

OAR and DWD are both supported by NINDS Tau Center without Walls Program (U54-NS100693) and NIH (UG3-NS104095). OAR is supported by NIH (P50-NS072187; R01-NS078086; U54-NS100693; U54-NS110435), DOD (W81XWH-17-1-0249) The Michael J. Fox Foundation, The Little Family Foundation, the Mayo Clinic Foundation, and the Center for Individualized Medicine. DWD receives research support from the NIH (P50-AG016574; U54-NS100693; P01-AG003949), CurePSP, the Tau Consortium, and the Robert E. Jacoby Professorship. ZKW is partially supported by the Mayo Clinic Center for Regenerative Medicine, the gifts from The Sol Goldman Charitable Trust, and the Donald G. and Jodi P. Heeringa Family, the Haworth Family Professorship in Neurodegenerative Diseases fund, and The Albertson Parkinson’s Research Foundation. SK is supported by a post-doctoral fellowship from the Karin & Sten Mortstedt CBD Solutions AB.

### Authors’ contributions

RRV designed the genotyping assays, performed all genotyping and quality control assessments, and drafted the manuscript. NT assisted with genotyping some samples. MGH and PWJ performed all statistical analysis and MGH provided manuscript improvements. AISB provided training for genotyping design and methods. RLW prepared DNA extracts for all samples from donated human tissues. SK and DWD provided brain tissue samples for all cases and DWD performed neuropathological assessments of PSP and CBD cases. RJU and ZKW recruited clinical patients and organised blood collections. OAR lead the study and oversaw all method developments and analysis and approved the final manuscript.

## Acknowledgements

We would like to thank the patients, donors, and caregivers who participated in this research, without who this work would not have been possible. We would like to acknowledge the continuous commitment, technical support and teamwork offered by Linda G. Rousseau, Virginia R. Phillips, and Monica Castanedes-Casey for tissue characterization. This work was supported in part by; the Mayo Clinic Florida Morris K. Udall Parkinson’s Disease Research Center of Excellence (NINDS P50 #NS072187); - an American Parkinson Disease Association (APDA), Mayo Clinic Information and Referral Center, and an APDA Center for Advanced Research. Samples included in this study were clinical controls or brain donors to the brain bank at Mayo Clinic in Jacksonville which is supported by CurePSP and the Tau Consortium. RRV designed the genotype assays and performed all laboratory analysis whilst MGH conducted, and is responsible for, all statistical data analysis. OAR had full access to all the data in the study and takes responsibility for the integrity of the data and the accuracy of the data analysis. The funding organizations and sponsors had no role in any of the following: design and conduct of the study; collection, management, analysis, and interpretation of the data; preparation, review, or approval of the manuscript; and decision to submit the manuscript for publication.

## Notes

**Financial Disclosures/Conflict of Interest**: None

