## Supplementary_data for "Associations of Mitochondrial Genomic Variation with Corticobasal Degeneration, Progressive Supranuclear Palsy, and Neuropathological Tau Measures"

### Supplementary Tables

| Neuroanatomical regions assessed for tau pathology | |
| --- | --- |
| Basal nucleus | Dentate nucleus |
| Caudate putamen | Inferior olive |
| Globus pallidus | Locus ceruleus |
| Hypothalamus | Medullary tegmentum |
| Motor cortex | Midbrain tectum |
| Subthalamic nucleus | Oculomotor complex |
| Temporal cortex | Pontine base |
| Thalamic fasciculus | Pontine tegmentum |
| Ventral thalamus | Red nucleus |
| Cerebellar white matter | Substantia nigra |

Supplementary Table 1: List of neuroanatomical regions assessed for tau pathology measures for neurofibrillary tangles (NFT), neuropil threads (NT), tufted astrocytes (TA), and oligodendroglial coiled bodies (CB) in N=767 PSP and N=152 CBD brains.

| **Assay Number** | **RSID** | **mtDNA Haplogroup Marker** | **Forward Primer Sequence** | **Reverse Primer Sequence** | **Unextended Primer Sequence** |
| --- | --- | --- | --- | --- | --- |
| 1 | rs879201732 | U1 | ACGTTGGATGGTGGGAAGAAGAAAGAGAGG | ACGTTGGATGACCAAATCTCCACCTCCATC | ACCTCCATCATCACCTC |
| 1 | rs28358280 | K | ACGTTGGATGTTGAGGGTTATGAGAGTAGC | ACGTTGGATGGGAATACTAGTATATCGCTC | TATCGCTCACACCTCAT |
| 1 | rs41473545 | H3 | ACGTTGGATGGGAGGTGAAATATGCTCGTG | ACGTTGGATGGCTTCCTAGGGTTTATCGTG | ATCGTGTGAGCACACCA |
| 1 | rs2853498 | U | ACGTTGGATGAGCTATCCATTGGTCTTAGG | ACGTTGGATGGGGTGGTTATAGTAGTGTGC | GGAGTTGCACCAAAATT |
| 1 | rs2298007 | A | ACGTTGGATGGGCCATGGCTAGGTTTATAG | ACGTTGGATGTTGCCACAACTAACCTCCTC | ATCCTCGGACTCCTGCCT |
| 1 | rs201950015 | J2 | ACGTTGGATGAAAGGAAGGAATCGAACCCC | ACGTTGGATGAAAGTCATGGAGGCCATGGG | GGGGTTGGCTTGAAACCA |
| 1 | rs193302980 | HV | ACGTTGGATGGGGAGGTCGATGAATGAGTG | ACGTTGGATGACAAGAACACCAATGACCCC | TTGACCCCAATACGCAAAA |
| 1 | rs2015062 | H | ACGTTGGATGATGATGGCAAATACAGCTCC | ACGTTGGATGGACATCGTACTACACGACAC | ACACGTACTACGTTGTAGC |
| 1 | rs869096886 | JT | ACGTTGGATGTTCTACACCCTAGTAGGCTC | ACGTTGGATGTGTTTAGTGAGCCTAGGGTG | GGGTGTTGTGAGTGTAAAT |
| 1 | rs2000975 | N | ACGTTGGATGTCAACAACCGACTAATCACC | ACGTTGGATGAGGTTCGTCCTTTAGTGTTG | TCCTTTAGTGTTGTGTATGG |
| 1 | rs2853503 | U5 | ACGTTGGATGAAGCGAGGTTGACCTGTTAG | ACGTTGGATGTTACTCTCATCGCTACCTCC | CGCCTATAGCACTCGAATAAT |
| 1 | rs28358585 | W | ACGTTGGATGTAGAAGAGCGATGGTGAGAG | ACGTTGGATGACCAAAGAGCCCCTAAAACC | TCCCCTAAAACCCGCCACATCT |
| 1 | rs2853495 | R0 | ACGTTGGATGGAGTGCGTTCGTAGTTTGAG | ACGTTGGATGCGCAGTCATTCTCATAATCG | GCATTCTCATAATCGCCCACGG |
| 1 | rs35788393 | HV0a | ACGTTGGATGGGTTTTCATCTCCGGTTTAC | ACGTTGGATGAAATGGGCCTGTCCTTGTAG | TGGCCTGTCCTTGTAGTATAAA |
| 1 | rs200336777 | J2b | ACGTTGGATGAGGACAACCAGTAAGCTACC | ACGTTGGATGGGGAGATAGTTGGTATTAGG | AGGATTGTTGTGAAGTATAGTA |
| 1 | rs2853826 | N | ACGTTGGATGTCTGGCCTATGAGTGACTAC | ACGTTGGATGGAGTCGAAATCATTCGTTTTG | GTTTAAACTATATACCAATTCGG |
| 1 | rs41423746 | U6 | ACGTTGGATGTTCGTTCGGTAAGCATTAGG | ACGTTGGATGACAACATACCCATGGCCAAC | TCTACTCCTCATTGTACCCATTCT |
| 1 | rs193302956 | R | ACGTTGGATGGGTTGTTAGCGGTAACTAAG | ACGTTGGATGCAGACCCAAACATTAATCAG | TCAGTTCTTCAAATATCTACTCAT |
| 1 | rs386828968 | N9 | ACGTTGGATGTGGTAAGGGCGATGAGTGTG | ACGTTGGATGACTCCCCATATCTAACAACG | GAACAACGTAAAAATAAAATGACA |
| 1 | rs2001030 | H2 | ACGTTGGATGCCAGAAAACTACGATAGCCC | ACGTTGGATGCCCTGTTCAACTAAGCACTC | TCTGTTCAACTAAGCACTCTACTCT |
| 1 | rs28358576 | U2'3'4'7'8'9' | ACGTTGGATGAGTACCGCAAGGGAAAGATG | ACGTTGGATGGCAGAAGGTATAGGGGTTAG | TTAGTCCTTGCTATATTATGCTTGG |
| 1 | rs878870695 | X | ACGTTGGATGCCGTAGACCTAACCATCTTC | ACGTTGGATGGTGATGAAATTGATGGCCCC | GGAAATTGATGGCCCCTAAGATAGA |
| 1 | rs3928306 | H1 | ACGTTGGATGCGAACCTTTAATAGCGGCTG | ACGTTGGATGTAGGGTTTACGACCTCGATG | ACCTCGATGTTGGATCAGGACATCCC |
| 1 | rs201361958 | R11'B6 | ACGTTGGATGAATTGGGCTGATTTGCCTGC | ACGTTGGATGAGACCCACAACAAATAGCCC | TAGACCCACAACAAATAGCCCTTCTAA |
| 2 | rs41419549 | H4 | ACGTTGGATGAGGGTTGTACGGTAGAACTG | ACGTTGGATGTAAACCAAACCCAGCTACGC | CCCAGCTACGCAAAATC |
| 2 | rs28358584 | U8b'c | ACGTTGGATGTACTACAACCCTTCGCTGAC | ACGTTGGATGGTAGAGGGTGATGGTAGATG | GCGGGTTTTAGGGGCTC |
| 2 | rs2248727 | N | ACGTTGGATGTTAGGAGTGGGACTTCTAGG | ACGTTGGATGTTTTACCACTCCAGCCTAGC | CTAGCCCCTACCCCCCAA |
| 2 | rs193302996 | P | ACGTTGGATGTTCGCCTACACAATTCTCCG | ACGTTGGATGGCTAGGATGAGGATGGATAG | TGCAAGGACGCCTCCTAG |
| 2 | rs3915611 | T2 | ACGTTGGATGGATTCGGGAGGATCCTATTG | ACGTTGGATGGTTCAACCAGTAACTACTAC | CTACTAATCAACGCCCATA |
| 2 | rs41479950 | T1 | ACGTTGGATGTACTTGCTTGTAAGCATGGG | ACGTTGGATGCCACCATGAATATTGTACGG | TGACCACCTGTAGTACATA |
| 2 | rs1057520074 | J2a | ACGTTGGATGACCAAATGCCCCTCATTTAC | ACGTTGGATGTAGGGAGGATATGAGGTGTG | GAAGTGAGATGGTAAATGC |
| 2 | rs527236203 | T | ACGTTGGATGGATCCTCCAAATCACCACAG | ACGTTGGATGTGGGCGATTGATGAAAAGGC | CAGGCGTCTGGTGAGTAGTG |
| 2 | rs41467651 | F | ACGTTGGATGGACTTAGGGCTAGGATGATG | ACGTTGGATGCCATGAGCCCTACAAACAAC | GGCCCTACAAACAACTAACCT |
| 2 | rs28357975 | V | ACGTTGGATGACCTGAGTAGGCCTAGAAAT | ACGTTGGATGACTTGATGGCAGCTTCTGTG | TAGAACTGGAATAAAAGCTAG |
| 2 | rs41347846 | I | ACGTTGGATGGTAAGGCTAGGAGGGTGTTG | ACGTTGGATGAATAGTACCGTTAACTTCC | CCGTTAACTTCCAATTAACTAG |
| 2 | rs28357681 | J1c | ACGTTGGATGAATACGCAAAACTAACCCCC | ACGTTGGATGCGAAGTTTCATCATGCGGAG | CATGGGGTGGGGAGGTCGATGA |
| 2 | rs28625645 | J | ACGTTGGATGTTAGCAGCGGTGTGTGTGTG | ACGTTGGATGTTATTTTCCCCTCCCACTCC | CTCCCATACTACTAATCTCATCAA |
| 2 | rs193302927 | N1 | ACGTTGGATGGGTAAAAGGAGGGCAATTTC | ACGTTGGATGCGCGTCCCTTTCTCCATAAA | CCATAAAATTCTTCTTAGTAGCTAT |
| 2 | rs879023568 | J1d | ACGTTGGATGTTGTCAACGTCAAGGAGTCG | ACGTTGGATGGGCGGACTAATCTTCAACTC | TTCAACTCCTACATACTTCCCCCATT |

Supplementary Table 2: Primer sequences for the two custom-designed iPlex arrays used to determine unique mitochondrial DNA haplogroup-defining SNPs in PSP, CBD, and control cohorts. Red text indicates failed SNPs.

|  | **Odds ratio we would have 80% power to detect at the P≤0.0019 significance level (associations with PSP) or the P≤0.0023 significance level (associations with CBD** | | **Effect size^1^ we would have 80% power to detect at the P≤0.0024 significance level (analysis of PSP cases) or the P≤0.0045 significance level (analysis of CBD cases** | |
| --- | --- | --- | --- | --- |
| **Haplogroup frequency** | **Associations with PSP  (vs. controls)** | **Associations with CBD  (vs. controls)** | **Associations with tau pathology scores in PSP cases** | **Associations with tau pathology scores in CBD cases** |
| 1% | 4.1 | 6.7 | 1.4 | 2.7 |
| 2% | 2.9 | 4.6 | 1.0 | 2.2 |
| 3% | 2.5 | 3.8 | 0.8 | 1.7 |
| 4% | 2.3 | 3.4 | 0.7 | 1.6 |
| 5% | 2.1 | 3.1 | 0.6 | 1.4 |
| 10% | 1.8 | 2.5 | 0.5 | 1.0 |
| 20% | 1.6 | 2.1 | 0.4 | 0.8 |
| 30% | 1.5 | 2.0 | 0.3 | 0.7 |
| 40% | 1.5 | 2.0 | 0.3 | 0.6 |
| 50% | 1.5 | 2.0 | 0.3 | 0.6 |

Supplementary Table 3: Odds ratios (associations with risk of PSP and CBD) and effect sizes (associations with tau pathology scores) that we had 80% power to detect in our study at the significance levels that we utilized after applying a Bonferroni correction for multiple testing. ^1^ Effect size is defined as the difference mean difference of the given tau pathology score between subjects with and without the given haplogroup divided by the standard deviation of the given tau pathology score. The P≤0.0019, P≤0.0023, P≤0.0045, and P≤0.0024 significance levels are those used in the analysis of our data for the given analysis after applying a Bonferroni correction for multiple testing. ^1^Effect size is defined as the difference mean difference of the given tau pathology score between subjects with and without the given haplogroup divided by the standard deviation of the given tau pathology score. P≤0.0019, P≤0.0023, P≤0.0045, and P≤0.0024 significance levels are those used in the analysis of our data for the given analysis after applying a Bonferroni correction for multiple testing.

|  | **Haplogroup frequency, No. (%)** | | | **PSP vs. controls** | | **CBD vs. controls** | |
| --- | --- | --- | --- | --- | --- | --- | --- |
| **Mitochondrial DNA Haplogroup** | **Controls (N=916)** | **PSP patients (N=1051)** | **CBD patients (N=173)** | **OR (95% CI)** | **P-value** | **OR (95% CI)** | **P-value** |
| NIWYAX | 67 (7.3%) | 77 (7.3%) | 11 (6.4%) | 0.97 (0.69, 1.37) | 0.86 | 0.78 (0.38, 1.47) | 0.47 |
| HV | 463 (50.5%) | 514 (48.9%) | 89 (51.4%) | 0.94 (0.78, 1.12) | 0.47 | 1.03 (0.74, 1.45) | 0.84 |
| JT | 172 (18.8%) | 230 (21.9%) | 40 (23.1%) | 1.25 (1.00, 1.56) | 0.052 | 1.42 (0.94, 2.11) | 0.088 |
| UK | 208 (22.7%) | 218 (20.7%) | 30 (17.3%) | 0.88 (0.71, 1.10) | 0.26 | 0.68 (0.43, 1.04) | 0.081 |

Supplementary Table 4: Associations of mitochondrial DNA super-haplogroups with risk of PSP and CBD (in comparison to controls) were evaluated using multivariable logistic regression models adjusted for age and sex. P-values ≤0.05 are considered statistically significant. OR=odds ratio; CI=confidence interval at 95%.

|  |  | **Association with CB tau pathology score** | | **Association with NFT tau pathology score** | | **Association with TA tau pathology score** | | **Association with NT tau pathology score** | |
| --- | --- | --- | --- | --- | --- | --- | --- | --- | --- |
| **Mitochondrial DNA Haplogroup** | **No. (%) in haplogroup (N=767)** | **Regression coefficient (95% CI)** | **P-value** | **Regression coefficient (95% CI)** | **P-value** | **Regression coefficient (95% CI)** | **P-value** | **Regression coefficient (95% CI)** | **P-value** |
| NIWYAX | 60 (7.8%) | -0.02 (-0.11, 0.07) | 0.61 | 0.02 (-0.06, 0.10) | 0.62 | -0.06 (-0.15, 0.04) | 0.23 | -0.03 (-0.12, 0.07) | 0.57 |
| HV | 366 (47.7%) | 0.01 (-0.04, 0.06) | 0.73 | 0.03 (-0.02, 0.07) | 0.25 | 0.00 (-0.05, 0.05) | 0.91 | 0.02 (-0.03, 0.07) | 0.42 |
| JT | 176 (22.9%) | -0.01 (-0.07, 0.05) | 0.68 | -0.05 (-0.10, 0.00) | 0.070 | 0.01 (-0.05, 0.07) | 0.70 | -0.06 (-0.12, 0.00) | 0.065 |
| UK | 160 (20.9%) | 0.00 (-0.06, 0.06) | 0.91 | 0.01 (-0.05, 0.06) | 0.83 | 0.01 (-0.05, 0.07) | 0.73 | 0.03 (-0.03, 0.10) | 0.31 |

Supplementary Table 5: Associations of mitochondrial DNA super-haplogroups with CB, NFT, TA, and NT tau pathology scores in PSP cases with measured tau pathology scores (N=767) from linear regression models that were adjusted for age at death, sex, Braak, and Thal phase. Regression coefficients are interpreted as the increase in mean CB, NFT, TA, or NT tau pathology scores for patients in the given mitochondrial DNA super-haplogroup compared to patients not in the given haplogroup (non-haplogroup). P-values ≤0.05 are considered statistically significant. CB=coiled bodies; NFT=neurofibrillary tangles; TA=tufted astrocytes; NT=neuropil threads; CI=confidence interval.

|  |  | **Association with CB tau pathology score** | | **Association with NFT tau pathology score** | | **Association with TA tau pathology score** | | **Association with NT tau pathology score** | |
| --- | --- | --- | --- | --- | --- | --- | --- | --- | --- |
| **Mitochondrial DNA Haplogroup** | **No. (%) in haplogroup (N=152)** | **Regression coefficient  (95% CI)** | **P-Value** | **Regression coefficient  (95% CI)** | **P-Value** | **Regression coefficient  (95% CI)** | **P-Value** | **Regression coefficient  (95% CI)** | **P-Value** |
| NIWYAX | 9 (5.9%) | -- | -- | -- | -- | -- | -- | -- | -- |
| HV | 76 (50.0%) | 0.02 (-0.07, 0.11) | 0.70 | 0.00 (-0.09, 0.08) | 0.95 | 0.00 (-0.04, 0.04) | 0.97 | -0.01 (-0.10, 0.09) | 0.91 |
| JT | 35 (23.0%) | 0.10 (-0.01, 0.20) | 0.071 | 0.02 (-0.08, 0.12) | 0.64 | 0.02 (-0.03, 0.07) | 0.37 | 0.06 (-0.06, 0.17) | 0.34 |
| UK | 29 (19.1%) | -0.14 (-0.25, -0.03) | 0.013 | 0.00 (-0.11, 0.10) | 0.94 | -0.04 (-0.09, 0.02) | 0.17 | -0.05 (-0.17, 0.07) | 0.43 |

Supplementary Table 6: Associations of mitochondrial DNA super-haplogroups with CB, NFT, TA, and NT tau pathology scores in CBD cases with measured tau pathology scores (N=152) from linear regression models that were adjusted for age at death, sex, Braak, and Thal phase. Regression coefficients are interpreted as the increase in mean CB, NFT, TA, or NT tau pathology scores for patients in the given mitochondrial DNA haplogroup compared to patients not in the given haplogroup (non-haplogroup). ^1^Statistical tests were not performed for these haplogroups owing to their rare frequency (<10 PSP cases in the given haplogroup). P-values ≤0.05 are considered statistically significant. CB=coiled bodies; NFT=neurofibrillary tangles; TA=tufted astrocytes; NT=neuropil threads; CI=confidence interval.

|  | | **Association with disease duration** | | **Association with age at onset** | |
| --- | --- | --- | --- | --- | --- |
| **Mitochondrial DNA**  **Haplogroup** | **No. (%) in haplogroup (N=440)** | **Regression Coefficient**  **(95% CI)** | **P-value** | **Regression Coefficient**  **(95% CI)** | **P-value** |
| N^1^ | 0 (0.0%) | -- | -- | -- | -- |
| N1^1^ | 3 (0.7%) | -- | -- | -- | -- |
| I | 10 (2.3%) | -0.89 (-2.88, 1.10) | 0.38 | -0.37 (-5.69, 4.96) | 0.89 |
| W | 11 (2.5%) | 2.81 (0.93, 4.68) | 0.003 | 0.20 (-4.86, 5.27) | 0.94 |
| N9^1^ | 1 (0.2%) | -- | -- | -- | -- |
| X^1^ | 6 (1.4%) | -- | -- | -- | -- |
| R and R0^1^ | 3 (0.7%) | -- | -- | -- | -- |
| HV and HV0a | 11 (2.5%) | -0.76 (-2.65, 1.14) | 0.43 | 2.34 (-2.72, 7.41) | 0.36 |
| H, H1, H2, H3 and H4 | 183 (41.6%) | -0.61 (-1.21, -0.01) | 0.045 | -0.67 (-2.27, 0.94) | 0.41 |
| H | 82 (18.6%) | -0.27 (-1.03, 0.49) | 0.49 | -2.05 (-4.07, -0.03) | 0.047 |
| H1 | 62 (14.1%) | -0.26 (-1.11, 0.59) | 0.55 | 0.17 (-2.10, 2.45) | 0.88 |
| H2 | 16 (3.6%) | -0.74 (-2.33, 0.84) | 0.36 | -1.35 (-5.59, 2.89) | 0.53 |
| H3 | 21 (4.8%) | -0.78 (-2.18, 0.61) | 0.27 | 3.64 (-0.06, 7.33) | 0.054 |
| H4^1^ | 2 (0.5%) | -- | -- | -- | -- |
| V | 15 (3.4%) | 2.05 (0.42, 3.67) | 0.014 | 0.89 (-3.48, 5.26) | 0.69 |
| JT^1^ | 0 (0.0%) | -- | -- | -- | -- |
| J, J1, J1d, J2a and J2b | 51 (11.6%) | 0.01 (-0.91, 0.94) | 0.98 | 1.91 (-0.55, 4.37) | 0.13 |
| J^1^ | 0 (0.0%) | -- | -- | -- | -- |
| J1 | 37 (8.4%) | -0.19 (-1.26, 0.88) | 0.73 | 1.62 (-1.22, 4.47) | 0.26 |
| J1d^1^ | 0 (0.0%) | -- | -- | -- | -- |
| J2a | 13 (3.0%) | 0.81 (-0.94, 2.56) | 0.36 | 2.89 (-1.77, 7.55) | 0.22 |
| J2b^1^ | 1 (0.2%) | -- | -- | -- | -- |
| T, T1 and T2 | 58 (13.2%) | 0.09 (-0.78, 0.97) | 0.84 | -1.40 (-3.73, 0.94) | 0.24 |
| T^1^ | 1 (0.2%) | -- | -- | -- | -- |
| T1 | 11 (2.5%) | 0.80 (-1.09, 2.70) | 0.40 | -0.64 (-5.71, 4.43) | 0.80 |
| T2 | 46 (10.5%) | -0.02 (-0.99, 0.95) | 0.97 | -1.24 (-3.83, 1.34) | 0.35 |
| U, U1, U3, U5, U6 and U8b'c | 62 (14.1%) | -0.14 (-0.99, 0.71) | 0.74 | -0.05 (-2.33, 2.22) | 0.96 |
| U | 26 (5.9%) | 0.44 (-0.81, 1.70) | 0.49 | -0.29 (-3.65, 3.06) | 0.86 |
| U1^1^ | 1 (0.2%) | -- | -- | -- | -- |
| U3^1^ | 0 (0.0%) | -- | -- | -- | -- |
| U5 | 35 (8.0%) | -0.45 (-1.54, 0.65) | 0.42 | 0.21 (-2.71, 3.13) | 0.89 |
| U6^1^ | 0 (0.0%) | -- | -- | -- | -- |
| K | 22 (5.0%) | 0.66 (-0.70, 2.02) | 0.34 | -0.12 (-3.77, 3.52) | 0.95 |
| NIWYAX | 35 (8.0%) | 0.98 (-0.11, 2.07) | 0.077 | 0.27 (-2.65, 3.19) | 0.86 |
| HV | 209 (47.5%) | -0.40 (-0.99, 0.19) | 0.18 | -0.30 (-1.89, 1.28) | 0.71 |
| JT | 109 (24.8%) | 0.06 (-0.62, 0.75) | 0.85 | 0.19 (-1.64, 2.03) | 0.84 |
| UK | 84 (19.1%) | 0.09 (-0.66, 0.84) | 0.81 | -0.08 (-2.09, 1.93) | 0.94 |

Supplementary Table 7: Associations of individual and super- mitochondrial DNA haplogroups with age of onset and disease duration in PSP cases for whom that information was available (N=440). For analysis of disease duration, regression coefficients, 95% CIs, and p-values result from linear regression models that were adjusted for age at PSP onset and sex. For analysis of age of PSP onset, regression coefficients, 95% CIs, and p-values result from linear regression models that were adjusted for sex. Regression coefficients are interpreted as the increase in mean disease duration or age of PSP onset for patients in the given haplogroup compared to patients not in the given haplogroup. P-values ≤0.0025 are considered statistically significant after applying a Bonferroni correction for multiple testing. ^1^Statistical tests were not performed for these haplogroups owing to their rare frequency (<10 PSP cases in the given haplogroup). CI=confidence interval.

### Supplementary Figures

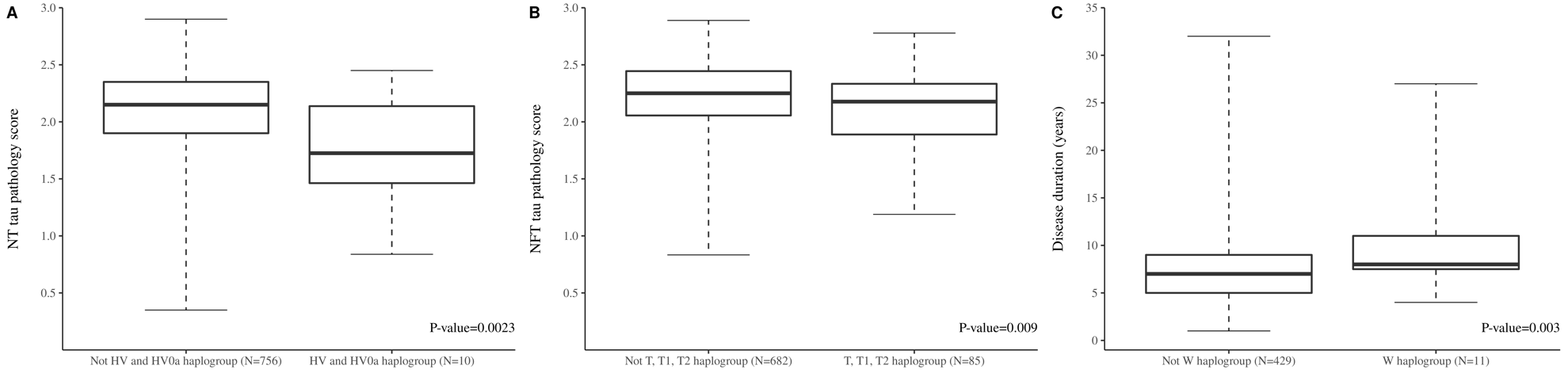

Supplementary Figure 1: Boxplots of [A] NT tau pathology scores between individuals with a HV and HV0a mtDNA haplogroup background compared to those who are not haplogroups HV/HV0a; [B] NFT tau pathology scores between individuals with a T, T1, and T2 mtDNA haplogroup background compared to those who are not haplogroups T/T1/T2; [C] difference in disease duration (years) between individuals with a mtDNA haplogroup W background compared to those who are not haplogroup W. NFT= neurofibrillary tangles; NT=neuropil threads.

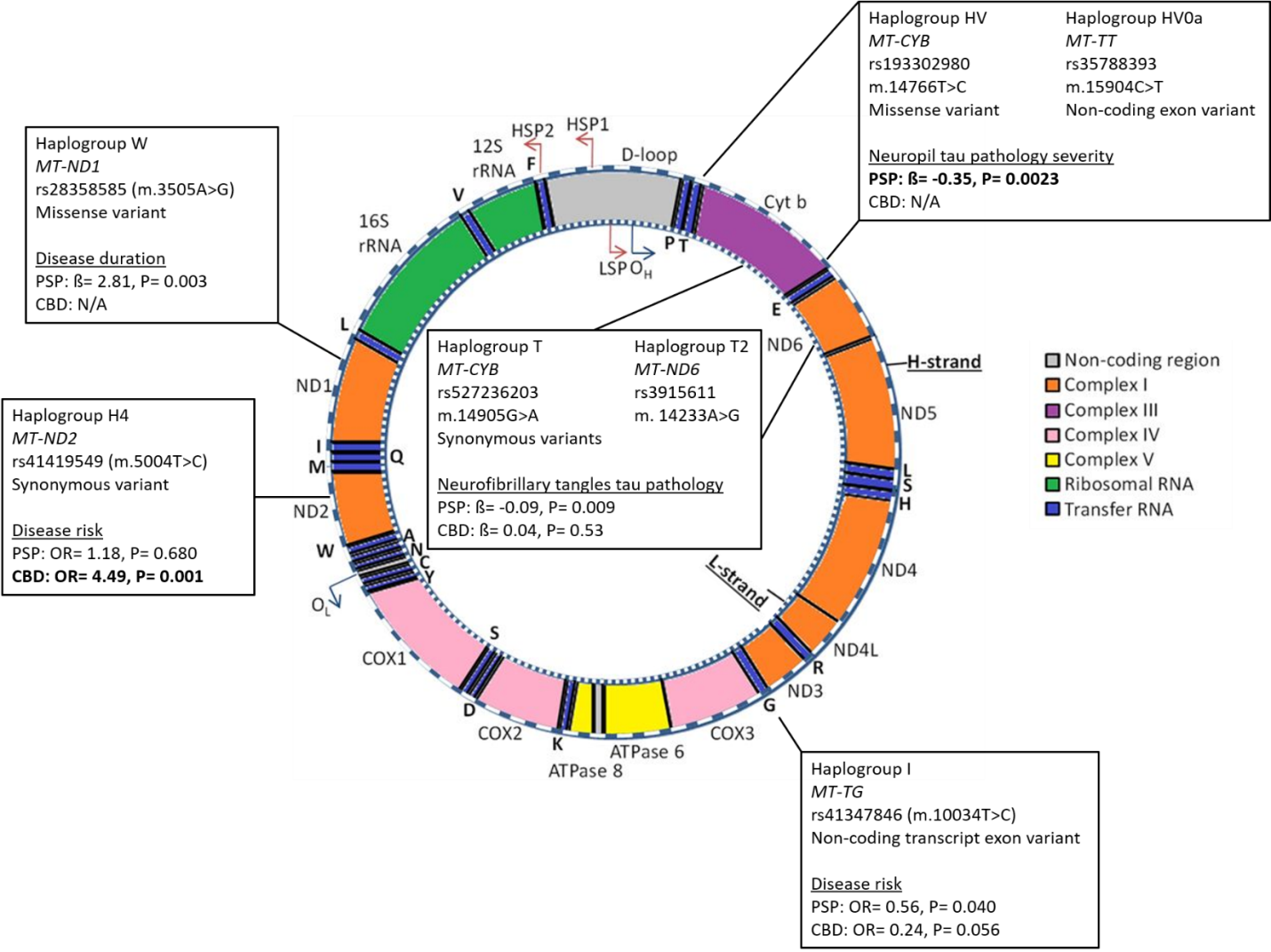

Supplementary Figure 2: Schematic diagram summarising significant results from this study. The diagram illustrates the circular 16,569bp mitochondrial genome and its respective genes. Protein-coding genes are coloured to indicate which subunit they code for and their respective complex in oxidative phosphorylation (orange= complex one; purple= complex three; pink= complex four; yellow= complex five). Transfer RNA (blue) letters indicate their respective amino acids and ribosomal RNA is highlighted in green. The D-loop region (grey) contains promoters for the heavy (H-) and light (L-) strands (HSP1, HSP2, and LSP respectively) and the origin of heavy-strand replication (OH). The origin of light-strand replication is indicated as OL. Text boxes detail the mitochondrial DNA haplogroup which was associated with which tested measurement in PSP and CBD cohorts, and details the unique SNP which was used to call for that specific haplogroup and the SNP location, RSID, mitochondrial DNA base position change, and variant type. This figure was adapted from Van der Wijst et al., 2017^1^

### Supplementary Methods

#### DNA Preparation and Genotyping

Unique haplogroup-defining mitochondrial DNA SNPs (mtSNPs) were selected from Phylotree^2^ and rsIDs determined from NCBI. DNA sequences were identified from the human NC012920.1 GRCh38 p.7 build (annotation 108). Sequences 200 base pairs surrounding each mtSNP were then generated from UCSC’s ‘blat’ tool, referencing the wildtype sequence, and custom primers were then manufactured. Two high multiplex iPlex assays (consisting of 39 mtSNPs - Figure 1) were automatically selected. Each genotyping chip was calibrated prior to sample measurements and contained eight or more randomly positioned negative replicates which were examined for contamination prior to determining samples genotypes. mtSNPs were excluded from analysis if present in >50% negative replicates, and the assay was accepted if individual negatives genotyped for <25% mtSNPs. rs2000975, rs28358576, rs193302996, and rs28357681, which called for haplogroups N, U2, P, and J1c respectively, failed and were removed from all analysis (≥90% success rate).

#### Mitochondrial DNA Haplogroup Assignment

Individuals who are mitochondrial DNA haplogroup H (i.e. not H1, H2, H3, or H4) would genotype for N, R, R0, HV, and H-defining variants, and did not genotype for additional H1, H2, H3, or H4 sub-haplogroup defining variants (Figure 1). For an individual to be further defined as a sub-haplogroup (e.g. H1, H2, H3, or H4) they would additionally need to genotype for their sub-haplogroup defining mtDNA SNPs as well as their main haplogroup variants (i.e all haplogroup H-defining mtDNA SNPs). Although the mitochondrial sub-haplogroup H1-defining variant rs3928306 defines other sub-haplogroup clades; to be defined H1, an individual would additionally have to have N, R, R0, HV, and H-defining variants. Mitochondrial super-haplogroups were defined by combining phylogenetic-related haplogroups together - e.g. individual haplogroups H and V (and their respective sub-haplogroups) were compiled to super-haplogroup HV.

1. van der Wijst MGP, van Tilburg AY, Ruiters MHJ, Rots MG. Experimental mitochondria-targeted DNA methylation identifies GpC methylation, not CpG methylation, as potential regulator of mitochondrial gene expression. Scientific Reports. 2017;7(1):177.
2. van Oven M, Kayser M. Updated comprehensive phylogenetic tree of global human mitochondrial DNA variation. Human Mutation. 2009;30(2):E386-E394.
